# Dopamine D1 and NMDA receptor co-regulation of protein translation in cultured nucleus accumbens neurons

**DOI:** 10.1101/2023.04.02.535293

**Authors:** Alexa R. Zimbelman, Benjamin Wong, Conor H. Murray, Marina E. Wolf, Michael T. Stefanik

## Abstract

Protein translation is essential for some forms of synaptic plasticity. We used nucleus accumbens (NAc) medium spiny neurons (MSN), co-cultured with cortical neurons to restore excitatory synapses, to examine whether dopamine modulates protein translation in NAc MSN. FUNCAT was used to measure translation in MSNs under basal conditions and after disinhibiting excitatory transmission using the GABA_A_ receptor antagonist bicuculline (2 hr). Under basal conditions, translation was not altered by the D1-class receptor (D1R) agonist SKF81297 or the D2-class receptor (D2R) agonist quinpirole. Bicuculline alone robustly increased translation. This was reversed by quinpirole but not SKF81297. It was also reversed by co-incubation with the D1R antagonist SCH23390, but not the D2R antagonist eticlopride, suggesting dopaminergic tone at D1Rs. This was surprising because no dopamine neurons are present. An alternative explanation is that bicuculline activates translation by increasing glutamate tone at NMDA receptors (NMDAR) within D1R/NMDAR heteromers, which have been described in other cell types. Supporting this, immunocytochemistry and proximity ligation assays revealed D1/NMDAR heteromers on NAc cells both *in vitro* and *in vivo*. Further, bicuculline’s effect was reversed to the same extent by SCH23390 alone, the NMDAR antagonist APV alone, or SCH23390+APV. These results suggest that: 1) excitatory synaptic transmission stimulates translation in NAc MSNs, 2) this is opposed when glutamate activates D1R/NMDAR heteromers, even in the absence of dopamine, and 3) antagonist occupation of D1Rs within the heteromers prevents their activation. Our study is the first to suggest a role for D2 receptors and D1R/NMDAR heteromers in regulating protein translation.

## INTRODUCTION

In many brain regions, dopamine (DA) and glutamate inputs converge onto common postsynaptic targets [1], with DA serving as a neuromodulator to influence synaptic transmission or plasticity at excitatory synapses [2–5]. Protein translation plays an important role in some forms of excitatory synaptic plasticity [6–9]. In hippocampus, signaling through D1 DA receptors (D1R) and protein kinase A (PKA) has long been implicated in the protein synthesis dependent phase of LTP [10–14], and D1Rs have also been shown to modulate protein translation directly [15,16].

Far less is known about the relationship between DA receptors and protein translation in the context of synaptic plasticity in medium spiny neurons (MSN) of the nucleus accumbens (NAc). These neurons play a critical role in motivated behaviors and disorders of motivation, including drug addiction [1,17]. It is now well established that synaptic plasticity plays a critical role in animal models of drug addiction [18–20]. More recently, dysregulation of protein translation has also been implicated in these models [21–31]. A fundamental question relevant to these lines of investigation is whether DA receptors modulate protein translation in the NAc.

Previously, to study other neuromodulatory processes in MSNs, we developed a co-culture system in which postnatal day 1 NAc neurons from rats are plated with prefrontal cortex (PFC) neurons from mice that express enhanced cyan fluorescent protein (ECFP) in all their cells; the cortical neurons restore excitatory synapses onto the MSNs, but the two cell types can be distinguished based on fluorescence [32]. Although there are no DA neurons in the co-cultures, MSNs express DA receptors and neuromodulatory effects of DA can be studied using exogenously applied agonists. For example, we demonstrated that activation of D1-class (D1/D5) DA receptors accelerates the insertion of GluA1-containing AMPARs onto the cell surface of MSNs and thereby facilitates their activity-dependent synaptic insertion [33,34,32]. More recently, using fluorescent noncanonical amino acid tagging (FUNCAT) in this co-culture system, we demonstrated that AMPA, NMDA and group I metabotropic glutamate receptors regulate translation in MSNs, in some cases in a manner distinct from what we observed in PFC neurons or what has been reported in hippocampus [35]. The purpose of the present study was to determine if DA receptors exert a neuromodulatory influence on protein translation in NAc MSNs in this co-culture system.

## MATERIALS AND METHODS

### Animals

All animal use procedures were approved by the Rosalind Franklin University of Medicine and Science Institutional Animal Care and Use Committee and were in accordance with the National Institutes of Health *Guide for the Care and Use of Laboratory Animals*. Pregnant Sprague Dawley rats (Harlan, Indianapolis, IN) were obtained at 18-20 days of gestation and housed individually in breeding cages. Postnatal day 1 (P1) offspring were decapitated and used to obtain NAc neurons. PFC neurons were obtained from P1 offspring of homozygous enhanced cyan fluorescent protein (ECFP)-expressing mice (strain: B6.129(ICR)-Tg(ACTB-ECFP)1Nagy/J; Jackson Laboratory, Bar Harbor, ME). This homozygous ECFP transgenic mouse strain was maintained in house by mating ECFP male and female mice. All offspring express ECFP.

### Postnatal nucleus accumbens/prefrontal cortex co-cultures

This NAc/PFC co-culture system has previously been described in detail [32,36]. Briefly, the medial PFC of ECFP-expressing P1 mice was dissected and dissociated with papain (20-25 U/mL; Worthington Biochemical, Lakewood, NJ) at 37°C for 30 minutes. The cells were then plated at a density of 30,000 cells/well onto coverslips coated with poly-D-lysine (0.1 mg/mL; Sigma, St. Louis, MO). One to three days later, the NAc from P1 rats was also dissected and dissociated with papain. These NAc cells were plated at a density of 30,000 cells/well with the PFC cells described previously. The NAc/PFC co-cultures were grown in Neurobasal media (Invitrogen, Carlsbad, CA), supplemented with 2 mM GlutaMAX, 0.5% Gentamicin, and 2% B27 (Invitrogen). Half of the media was replaced every 4 days. Cultures were used for experiments between 14 and 21 days in vitro.

## FUNCAT

Direct detection of de novo protein synthesis was achieved using a strategy adapted from previous studies [37–39] based on incorporation of the non-canonical amino acid L-azidohomoalanine (AHA) into newly translated proteins and subsequent fluorescent tagging of AHA-bearing proteins using click chemistry. Co-cultures were grown for 14-21 days in supplemented Neurobasal media (see previous section) until the day of the experiment when media was replaced with supplemented methionine-free DMEM (Invitrogen) for 30 minutes. This methionine (Met) starvation enhances labeling by AHA, which incorporates at methionine codons. After Met starvation, AHA (1 mM, pH 7.4; Life Technologies) was added to the wells (with or without test drugs) for 2 h. In addition, every experiment included a control group (4-6 wells) in which, after Met starvation, DMEM supplemented with 1 mM Met was added for 2 h. This group was used to define the background signal in the absence of AHA incorporation for each experiment (see Image Quantification and Statistical Analysis, below). After AHA or Met incubation, cells (on coverslips) were washed, fixed in 4% PFA for 15 minutes, and stored in PBS until click chemistry reactions. Cells were permeabilized with 0.25% Triton X-100, washed with 3% BSA, and the click chemistry reaction was performed by incubating the coverslips upside-down on droplets of PBS, pH 7.8, containing a Cy-5 conjugated copper-free click chemistry reagent, dibenzocyclooctyne (DBCO, 20nM, Click Chemistry Tools, Scottsdale, AZ) for 30 minutes. Coverslips were then mounted on slides using ProLong Gold Antifade mountant (ThermoFisher, Waltham, MA).

### Drug treatments

To determine their effect on protein translation, the following drugs were incubated for 2 h along with AHA as described above: R(+)-SKF-81297 hydrobromide (SKF, 1 μM, Sigma-Aldrich, St. Louis, MO); (4a*R*-trans)-4,4a,5,6,7,8,8a,9-Octahydro-5-propyl-1*H*-pyrazolo[3,4-g]quinolone (quinpirole, 1 μM, Sigma-Aldrich); [R-(R*,S*)]-6-(5,6,7,8-Tetrahydro-6-methyl-1,3-dioxolo[4,5-g]isoquinolin-5-yl)furo[3,4-e]-1,3-benzodioxol-8(6*H*)-one (bicuculline, 20 μM, Tocris); SCH23390 hydrochloride (SCH23390, 10 μM, Tocris); 3-Chloro-5-ethyl-*N*-[(2*S*)-1-ethyl-2-pyrrolidinyl)methyl]-6-hydroxy-2-methoxybenzamide (Eticlopride, 1 μM, Tocris); D-(-)-2-Amino-5-phosphonopentanoic acid (APV,50 μM, Tocris).

### Immunocytochemistry

After 14-21 days in vitro, NAc/PFC co-cultures were washed with PBS and fixed in 4% PFA for 20 minutes, washed again, and stored in PBS until the experiment. To perform the immunocytochemistry, the fixed cells were washed in 1X PBS, permeabilized in PBS containing 0.1% Triton X-100 for 15 minutes and washed again, before being blocked for 1 hr in 3% bovine serum albumin (BSA) in PBS, pH 7.4, at room temperature. Cells were then incubated with one of the following primary antibodies overnight at 4°C: 1:2000 rat anti-D1R (D2944, Sigma Aldrich; gift of Susan George), 1:1000 rabbit anti-GluN1 (D65B7 #5704S; Cell Signaling Technology), or 1:1000 mouse anti-tyrosine hydroxylase (#22941; Immunostar, Hudson, WI). Coverslips were then washed with PBS and incubated in 1:500 Cy3-conjugated rabbit anti-mouse (PA43009V, GE Healthcare, Buckinghamshire, UK), 1:500 Cy3-conjugated goat anti-rabbit (28901106V, GE Healthcare) or goat anti-rat (A10525, Invitrogen, Carlsbad, CA) secondary antibody for 1 hr at room temperature before being mounted on slides using ProLong Gold Antifade mountant (Thermo Fisher, Waltham, MA).

For GluA1/GluA2 immunohistochemical slice experiments, rats were anesthetized with chloral hydrate (16% w/v in sterile saline, Sigma-Aldrich), and then transcardially perfused with PBS, followed by PBS containing 4% w/v paraformaldehyde. Brains were postfixed for 24 hr at room temperature in perfusion solution. Coronal sections (30 µm thick) were incubated for 1 hr in 5% normal donkey serum (NDS) and 0.3% Triton X-100, and then incubated overnight in 5% NDS plus primary antibodies against GluA1 (1:1000, ms2263, Millipore) and GluA2 (1:1000, rb53065, Cell Signaling Technology). Sections were rinsed 3x in PBS, incubated for 1 hr in 1:500 dilutions of the secondary antibodies (Cy5-conjugated goat anti-mouse, PA45009V, GE Healthcare; Cy3-conjugated goat anti-rabbit, #2890116V, GE Healthcare), and then rinsed 3x in PBS before mounting on slides using ProLong Gold Antifade mountant.

### Proximity ligation assay

The proximity ligation assay (PLA) protocol was performed as described by the manufacturer (Duolink, Sigma-Olink). Briefly, this assay detects two epitopes that are in very close proximity, i.e., in the same molecule or heteromer [40–42]. Standard primary antibodies to the two epitopes are used, along with a pair of oligonucleotide-labeled secondary antibodies (PLA probes) included in the kit. If the two target epitopes and therefore the two PLA probes are in close proximity, linker oligonucleotides and a ligase promote formation of a ‘circle’ subsequently amplified by rolling-circle amplification and visualized by fluorescently labeled probes. In our study, coverslips containing co-cultured neurons (described above) or 30 µm coronal slices from the rat brain were incubated for 1 hr at 37°C with the blocking solution in a preheated humidity chamber, followed by an overnight incubation with either the primary D1R and GluN1 antibodies or the GluA1 and GluA2 antibodies described above (rat anti-D1R at 1:400 and rabbit anti-GluN1 at 1:50; or mouse anti-GluA1 at 1:1000 and rabbit anti-GluA2 at 1:1000). To allow for the detection of the rat anti-D1R antibody by the anti-mouse PLA probe, following the primary antibody incubation, samples were washed and then again incubated overnight with a mouse anti-rat secondary antibody (1:500; 13-4813-85, Invitrogen). The next day, cells or slices were washed with buffer A (DUO82047, Sigma-Olink), and incubated for 1 h with the Duolink In Situ PLA anti-rabbit PLUS probe (DUO9002, Sigma-Olink) and the anti-mouse MINUS probe (DUO92004, Sigma-Olink). The PLA signal was detected using the Duolink in situ PLA detection kit (DUO92007, Sigma-Olink). Nuclei were labeled by a DAPI mounting media included with the labeling kit. Positive PLA signals were visualized as red dots. Experiments utilizing the D1R antibody were also conducted by conjugating the PLA MINUS probe directly to the antibody using the Duolink In Situ Probemaker kid (DUO92010, Sigma-Olink). There was no statistically significant difference in puncta number using this approach as compared to using anti-mouse detection of the anti-rat antibody (t(12)= 0.2335, p = 0.3846). Data presented in figures were generated using the latter method. Negative control assays were performed to ensure the specificity of the PLA labeling and amplification. These included experiments carried out in the absence of one of the two PLA probes, or in the absence of the ligase and polymerase. No PLA signal was observed in these conditions (data not shown). Negative control data showing no PLA signal without the D1R primary antibody are shown in Figure 7. As a positive control, we used PLA to detect GluA1/GluA2 heteromeric AMPARs on processes of MSN in our co-culture system (see Results for additional discussion).

### Image quantification and statistical analysis

Images were acquired and analyzed with an imaging system consisting of a Nikon (Melville, NY) inverted microscope, ORCA-ER digital camera, and MetaMorph software (Universal Imaging, Downingtown, PA). Images for all experimental groups were taken using identical acquisition parameters. All groups to be compared were processed simultaneously using cells from the same culture preparation. For each experimental group, cells from at least 4 different wells were used, and approximately 3-6 cells from each well were analyzed. We restricted our analysis to NAc MSNs, which can be distinguished from other neuronal cell types based on morphology and from ECFP-expressing PFC cells based on the absence of fluorescent signal [32]. MSNs were selected for analysis under phase contrast imaging to avoid experimenter bias and images were coded to ensure analysis was blind to experimental condition. For FUNCAT experiments, AHA labeling was quantified as the total area of Cy5 fluorescence in a fixed length (15 μm) of a process located at least one soma diameter away from the soma. We have previously demonstrated modulatory effects of DA on AMPAR trafficking in MSN processes selected according to these parameters [33,34,32,36]. The background threshold was set based on an average of background fluorescence in unstained areas in negative control wells (incubated with Met but not AHA; see above). Met wells were included in ANOVA calculations, however are not included in graphs because fluorescence levels were extremely low compared to other groups. Values for drug + AHA cells were normalized to the mean of the AHA-only group. For receptor quantification experiments, images were acquired using identical acquisition parameters for each receptor. Neuronal processes were selected for analyses under phase contrast imaging and fluorescently labeled D1R and GluN1 were measured using a threshold set at least two times higher than the average background fluorescence. D1R expression alone was defined as punctate staining that did not overlap with GluN1, and vice versa. Quantification of puncta in brain slices was conducted using ImageJ software (NIH). Images were thresholded to distinguish signal from background; the same threshold was applied to all images. The number of puncta were quantified using the “Analyze Particles” macro with the exclusion criteria of size of objects set at greater than 5 µm^2^. Experimental data were analyzed for significance with one-way ANOVAs followed by Tukey’s multiple comparison tests (Figures 1 and 2) or unpaired t-test (Figure 7), using Prism 6 (Graphpad) software. Statistical significance was defined as p<0.05 and all data are presented as mean +/-SEM.

**Figure 1.**
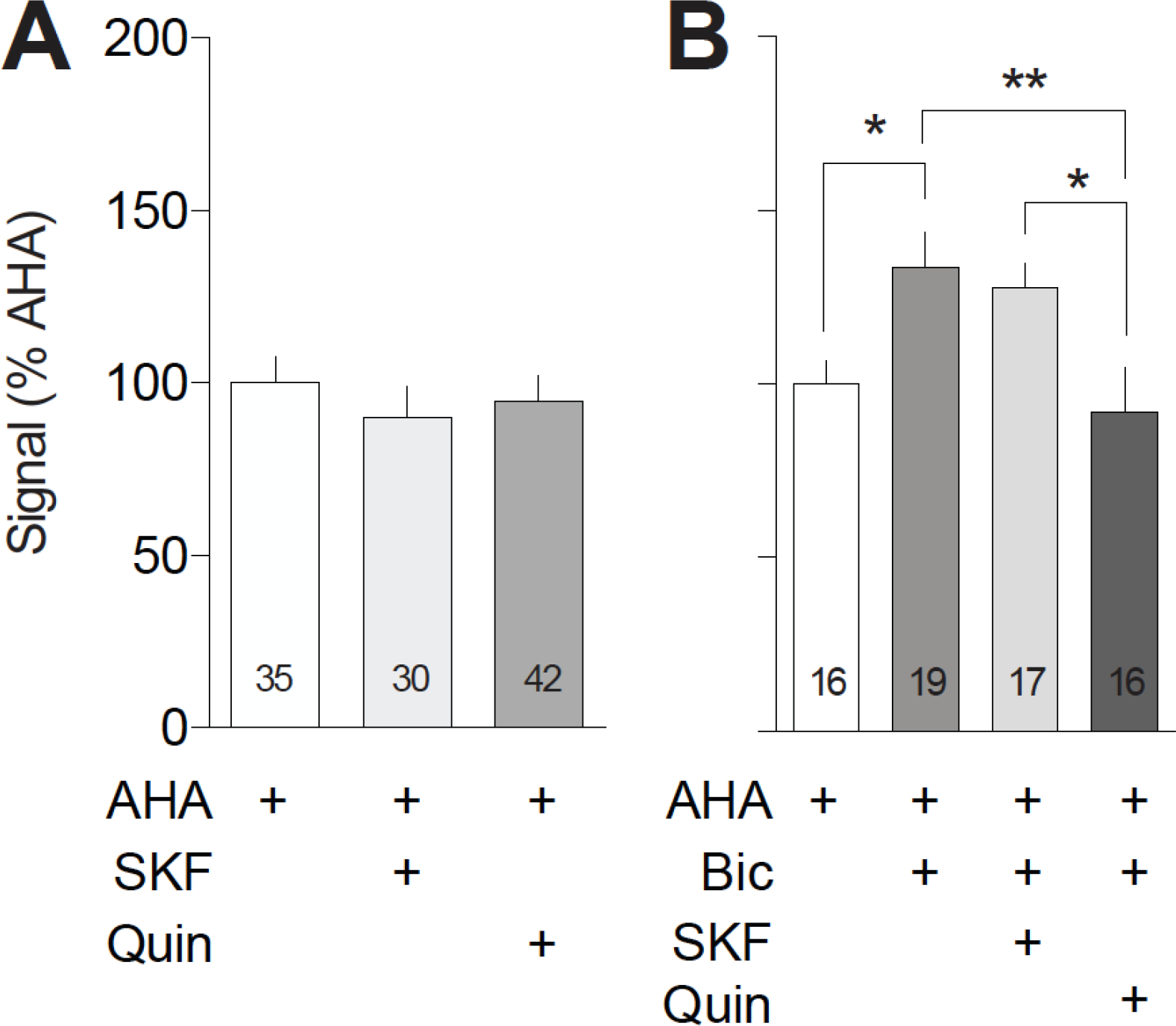
Dopamine agonists fail to alter basal translation in cultured NAc neurons but a D2-class agonist reverses bicuculline-stimulated translation. Left: Co-cultured NAc and PFC neurons were incubated with 1 mM AHA plus or minus the dopamine D1-class agonist SKF81297 (SKF; 1 µM) or the D2-class agonist quinpirole (Quin; 50 µM) for 2 h and tagged with 20 nM DBCO-Cy5. The addition of either SKF or Quin failed to alter protein translation in the processes of NAc MSNs [F(2, 104)=0.23; p=0.69]. The number of processes analyzed is indicated within the bars in this and subsequent figures. Right: Co-cultured NAc and PFC neurons were incubated with 1 mM AHA plus or minus the GABA_A_ receptor antagonist bicuculline (Bic; 20 µM) and/or SKF81297 (SKF; 1 µm) or quinpirole (Quin; 50 µM) for 2 h and tagged with 20 nM DBCO-Cy5. Bicuculline increased translation in the processes of NAc MSNs [F(4,80)=11.22, p<0.0001; Tukey’s post hoc p=0.04]. This effect was not impacted by co-incubation with SKF81297 (Tukey’s post hoc p=0.984), but was significantly reduced by quinpirole (Tukey’s post hoc p=0.036). *p<0.05, **p<0.005

**Figure 2.**
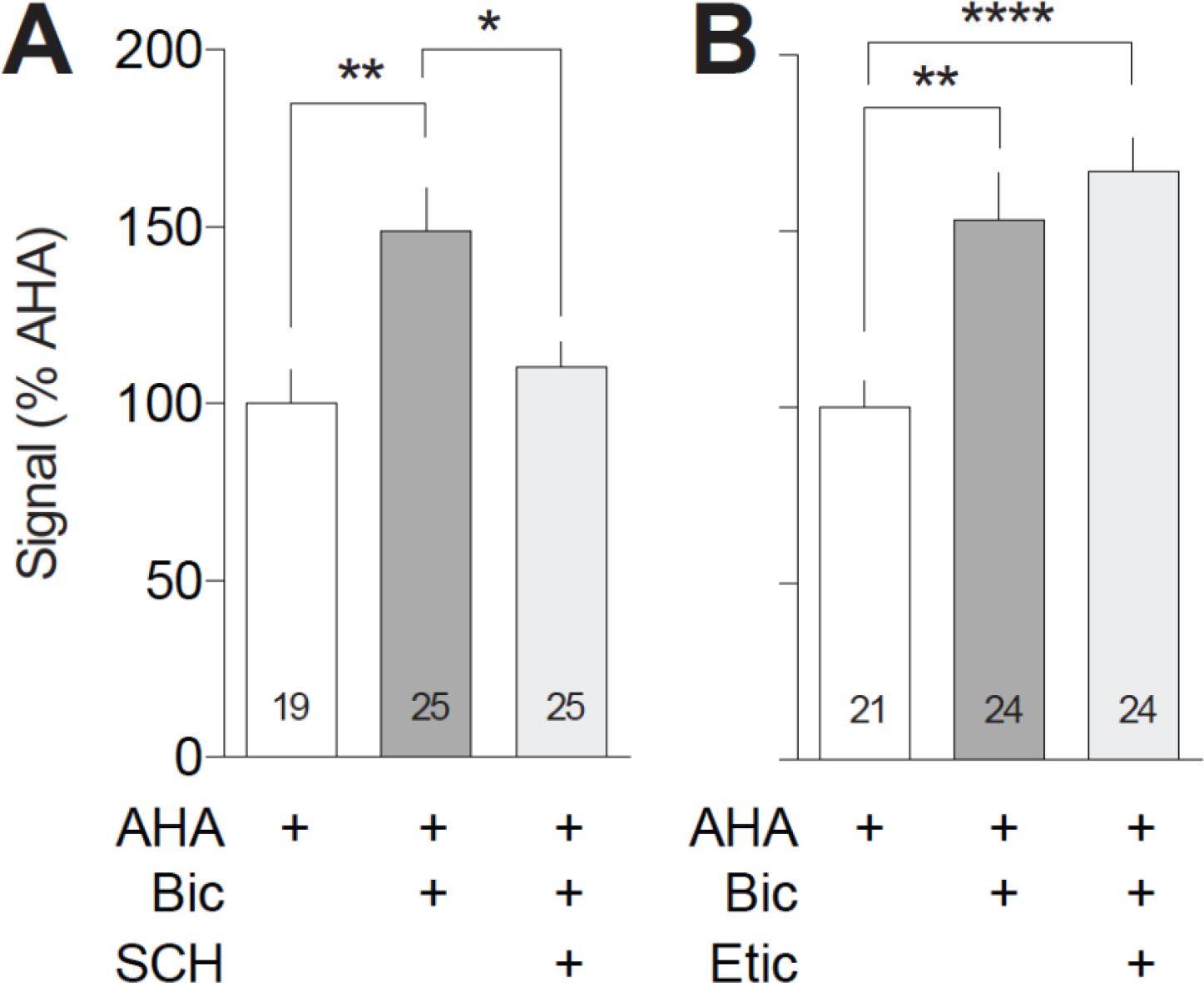
Bicuculline-stimulated translation is unaffected by a D2-class antagonist but is blocked by a D1-class antagonist. Left: Co-cultured NAc and PFC neurons were incubated with 1 mM AHA plus or minus the GABA_A_ receptor antagonist bicuculline (Bic; 20 µM) and/or the dopamine D1-class antagonist SCH23390 (SCH; 1 µM) for 2 h and tagged with 20 nM DBCO-Cy5. Bicuculline increased translation in the processes of NAc MSNs [F(2,66)=1.40, p=0.003; Tukey’s post hoc p=0.004]. This effect was blocked by SCH23390 (Tukey’s post hoc p=0.018). Right: Co-cultured NAc and PFC neurons were incubated with 1 mM AHA plus or minus the GABA_A_ receptor antagonist bicuculline (Bic; 20 µM) and/or the dopamine D2-class antagonist eticlopride (Etic; 1 µM) for 2 h and tagged with 20 nM DBCO-Cy5. Bicuculline increased translation in the processes of NAc MSNs [F(3,75)=10.93; p<0.0001, Tukey’s post hoc p=0.002]. This effect was not impacted by co-incubation with eticlopride (Tukey’s post hoc p=0.754). *p<0.05, **p<0.005, ****p<0.0001

## RESULTS

### FUNCAT measures of translation in cultured NAc MSNs

In order to visualize newly synthesized proteins in cultured NAc MSNs, we utilized FUNCAT. This technique employs a non-canonical amino acid bearing an azide group (azidohomoalanine; AHA), which incorporates into newly synthesized proteins at methionine codons [43,37,44]. Proteins that have been newly synthesized and therefore incorporated AHA can then be labeled using click chemistry in copper-free conditions with a fluorescent reporter containing dibenzylcyclooctyne (DBCO) [45,46,43].

We have previously validated FUNCAT as a measure of protein translation in processes of NAc MSNs in the same co-culture system utilized here [35]. Briefly, we showed that AHA incorporation was abolished by the protein translation inhibitor cycloheximide and colocalizes with the ribosomal protein S6, a marker of translation. Furthermore, AHA labeling in processes was adjacent to synapses, consistent with regulation of dendritic translation by synaptic activity [47,48]. Based on our prior results [35], all cultures in the present study were incubated with AHA for 2 h, in the presence or absence of test drugs, as this duration of AHA labeling provides a strong and reproducible signal.

### DA receptor agonists fail to alter basal translation

Previous work has implicated DA in regulating protein translation in hippocampus [15,16]. We have previously demonstrated the existence of D1, D5 and D2 receptors on NAc MSNs in our co-culture system [32]. Here we used application of exogenous DA receptor-selective drugs to determine whether DA receptors modulate translation in NAc MSNs (Fig. 1). From this point forward, we will use the term D1-class to refer to the D1 family of receptors (which includes the D5 receptor) because agonists and antagonists used in our studies act on both D1 and D5 receptors, and adopt a similar terminology for the D2 family (which includes D2, D3 and D4 receptors). Co-cultures were incubated for 2 h with AHA along with either the D1-class receptor agonist SKF81297 (1 µM) or the D2-class receptor agonist quinpirole (1 µM). These drug concentrations have previously been shown to be sufficient to modulate AMPAR trafficking in the present co-culture system and in cultured cortical neurons [32,49]. In this and all subsequent experiments, co-cultures treated with AHA + drugs were always compared to a control group treated with AHA only; this group provides a measure of basal translation. Fig. 1A shows that neither SKF81297 nor quinpirole impacted basal translation relative to the AHA-only control group.

### A D1-class, but not D2-class, receptor antagonist attenuates bicuculline-stimulated protein translation

We showed recently that incubating NAc/PFC co-cultures with bicuculline is sufficient to increase protein translation in NAc MSNs, and that this increase is due to glutamate receptor signaling, as it can be blocked by co-administration of glutamate receptor antagonists [35]. Therefore, we next assessed whether DA receptors modulate bicuculline-stimulated protein translation in NAc MSNs.

We began by incubating cells with bicuculline plus either the D1-class agonist SKF81297 (1 µM) or the D2-class agonist quinpirole (1 µM) for 2 h. Quinpirole but not SKF81297 significantly reduced the bicuculline-stimulated increase in protein translation to the level observed in control cells incubated with AHA alone (Fig. 1B). Next, we treated co-cultures with bicuculline plus either the D1-class antagonist SCH23390 (1 µM) or the D2-class antagonist eticlopride (1 µM) for 2 h. These drug concentrations were selected based on previous studies in this and other culture systems [50,32]. We observed that bicuculline-stimulated translation was reduced by SCH23390 but was unaffected by eticlopride (Fig. 2). These results indicate that bicuculline-stimulated protein translation is reduced by a D2-class agonist or a D1-class antagonist.

### Evidence for functional D1/NMDAR heteromers in NAc MSNs

Reduction of bicuculline-driven translation by a D1-class antagonist is surprising, as cells capable of producing DA are not present in either the PFC or NAc, and thus should not be present in the co-cultures. We confirmed there were no DA neurons in these cultures by staining for the dopaminergic marker, tyrosine hydroxylase (TH; Fig. 3). Therefore, although we have previously shown that both D1-class (D1 and D5) and D2 receptors are expressed by MSNs in the co-culture system [32], there should not be endogenous dopaminergic tone at these receptors. One possible explanation for nevertheless observing an effect of SCH23390 is that D1-class receptors are interacting with glutamate receptors to produce the observed reduction in translation. Indeed, multiple techniques have confirmed the existence of D1R and NMDA receptor (NMDAR) heteromers in brain regions including striatum (see Discussion). As a first step to test this hypothesis, we evaluated potential colocalization of D1Rs and NMDARs in NAc MSNs using immunocytochemistry. For these experiments, we selectively analyzed NAc MSNs (PFC neurons were excluded based on their cyan fluorescence; see Methods). Fig. 4 shows a representative image of a NAc MSN process demonstrating puncta positive for the obligatory NMDAR subunit GluN1 (green) or the D1R (red), as well as a small percentage of puncta (∼15%, yellow) co-localizing GluN1 and the D1R. These results are consistent with the presence of D1R/NMDAR heteromers in MSNs in our co-culture system.

**Figure 3.**
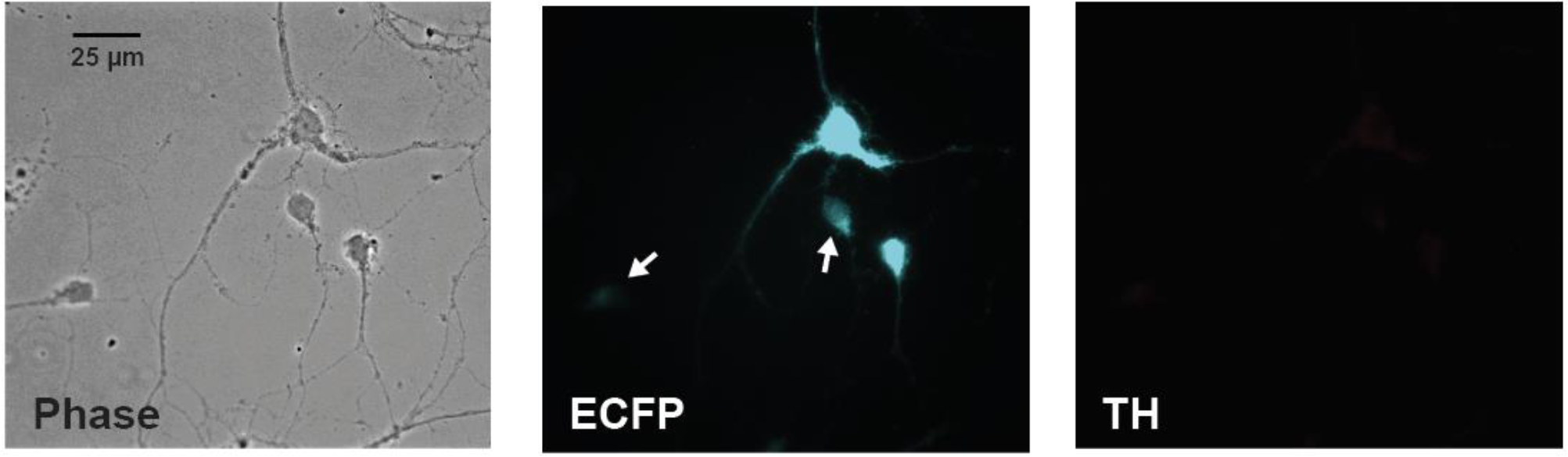
NAc/PFC co-cultures do not contain dopaminergic neurons. Representative images show that neurons in the NAc/PFC co-culture system do not express tyrosine hydroxylase (TH; right panel), a marker of dopamine neurons. PFC neurons are positive for cyan fluorescent protein (ECFP) because they are prepared from mice expressing ECFP in all their cells. Arrows indicate NAc neurons, which exhibit only low autofluorescence.

**Figure 4.**
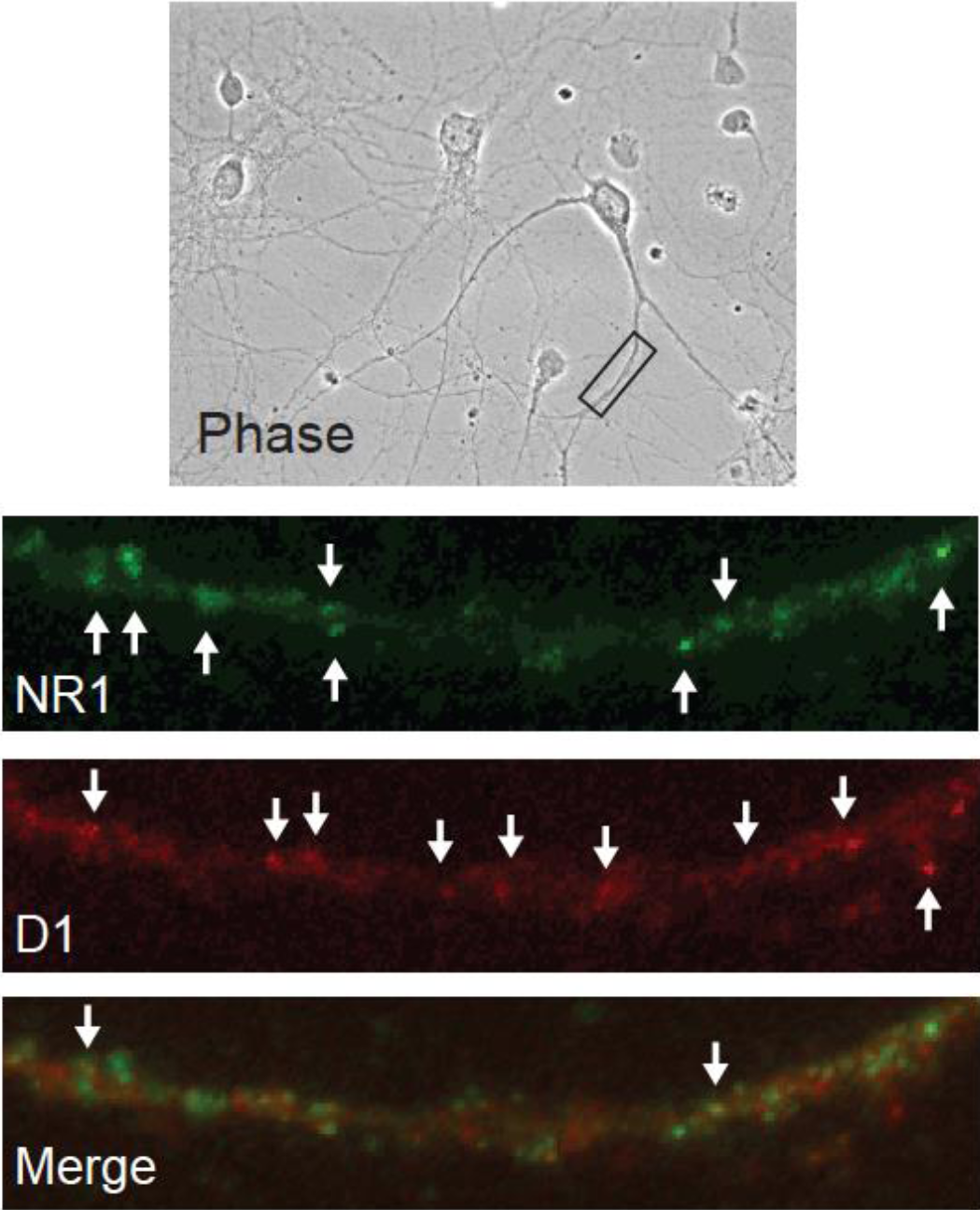
Immunocytochemistry reveals co-localization of the D1 receptor and GluN1 in cultured NAc neurons. Representative images indicate the presence of the obligatory NMDA receptor subunit GluN1 (green), the dopamine D1 receptor (red) and a small population of puncta in which GluN1 and D1 receptors are co-localized (merge), consistent with close proximity of the receptors in a heteromeric complex.

To provide direct evidence of D1R/NMDAR heteromers, we utilized a proximity ligation assay (PLA) in cultured neurons and in brain slices from the NAc of adult rats. This technique, which generates a signal if two epitopes are in extremely close proximity, has previously been used to show receptor-receptor interactions in both cultured neurons and brain slices [51,52,40]. Control experiments were performed to validate the specificity of this signal. No PLA puncta were observed in the absence of one of the two PLA probes, or in the absence of the ligase and polymerase (data not shown). As a positive control, we used PLA to detect GluA1/GluA2 heteromeric AMPARs on processes of MSN in our co-culture system and in brain slices taken from the NAc (Fig. 5). This is the most common AMPAR subtype in principal neurons [53–55]. We have previously demonstrated their presence on co-cultured NAc MSNs using immunocytochemistry [36] and in the NAc of intact rats using co-immunoprecipitation [56,54]. Having validated the assay, we used PLA to test for D1R/GluN1 heteromers. In agreement with the immunocytochemical data (Fig. 4), we observed PLA signals (red puncta) on the processes of cultured NAc MSNs (Fig. 6), suggesting a close proximity of the D1R and GluN1 in this cell type. Next, we sought to determine whether or not these heteromers exist within NAc neurons of the adult rat brain by conducting PLA assays on coronal brain sections (30 µm). PLA signals, indicating the presence of D1R/NMDAR heteromers, were observed in the proximity of NAc cell bodies identified by DAPI staining (Fig. 7). These results are in agreement with our observations in cultured NAc neurons.

**Figure 5.**
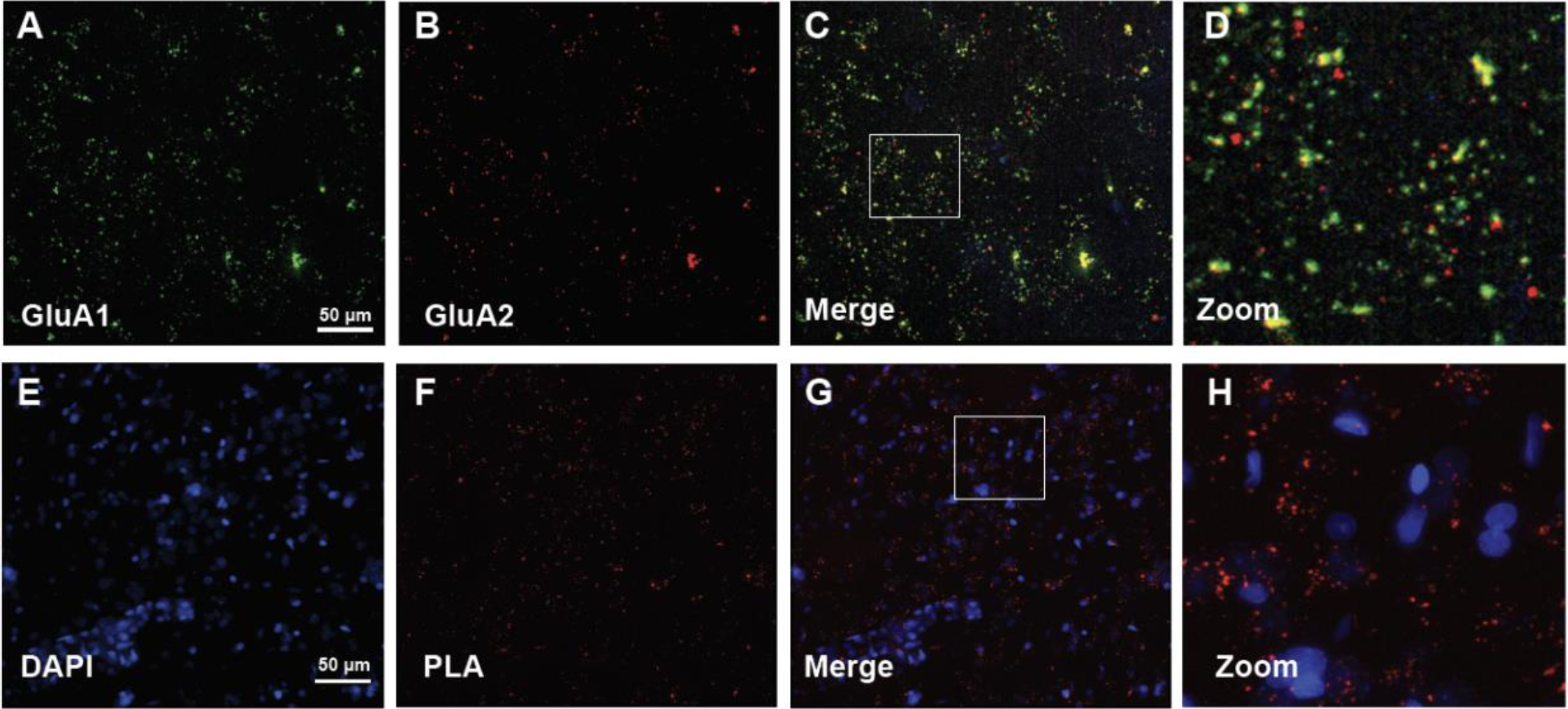
Demonstration of GluA1/GluA2-containing AMPARs in brain slices using a proximity ligation assay. GluA1 and GluA2 were analyzed using both traditional co-localization of fluorescent imaging and a proximity ligation assay (PLA) in NAc medium spiny neurons (MSNs). (A) Fluorescently-labeled GluA1-containing AMPA receptors. (B) Fluorescently-labeled GluA2-containing AMPA receptors. (C, D) Immunocytochemistry reveals GluA1/GluA2 colocalization in NAc MSNs in rat brain slices (yellow puncta in merged image). This was expected because GluA1/GluA2-containing AMPA receptors are common in MSNs (see Results). Bottom Row: GluA1 and GluA2 were analyzed using PLA. PLA signals (red dots) indicate the presence of GluA1/GluA2-containing AMPARs. Representative images show neuronal nuclei (stained by DAPI; E), PLA signals (red dots; F), and merged DAPI and PLA images (G) demonstrating GluA1/GluA2 heteromers in proximity to NAc cell bodies (box indicates the area shown at higher magnification in the far-right panel; H).

**Figure 6.**
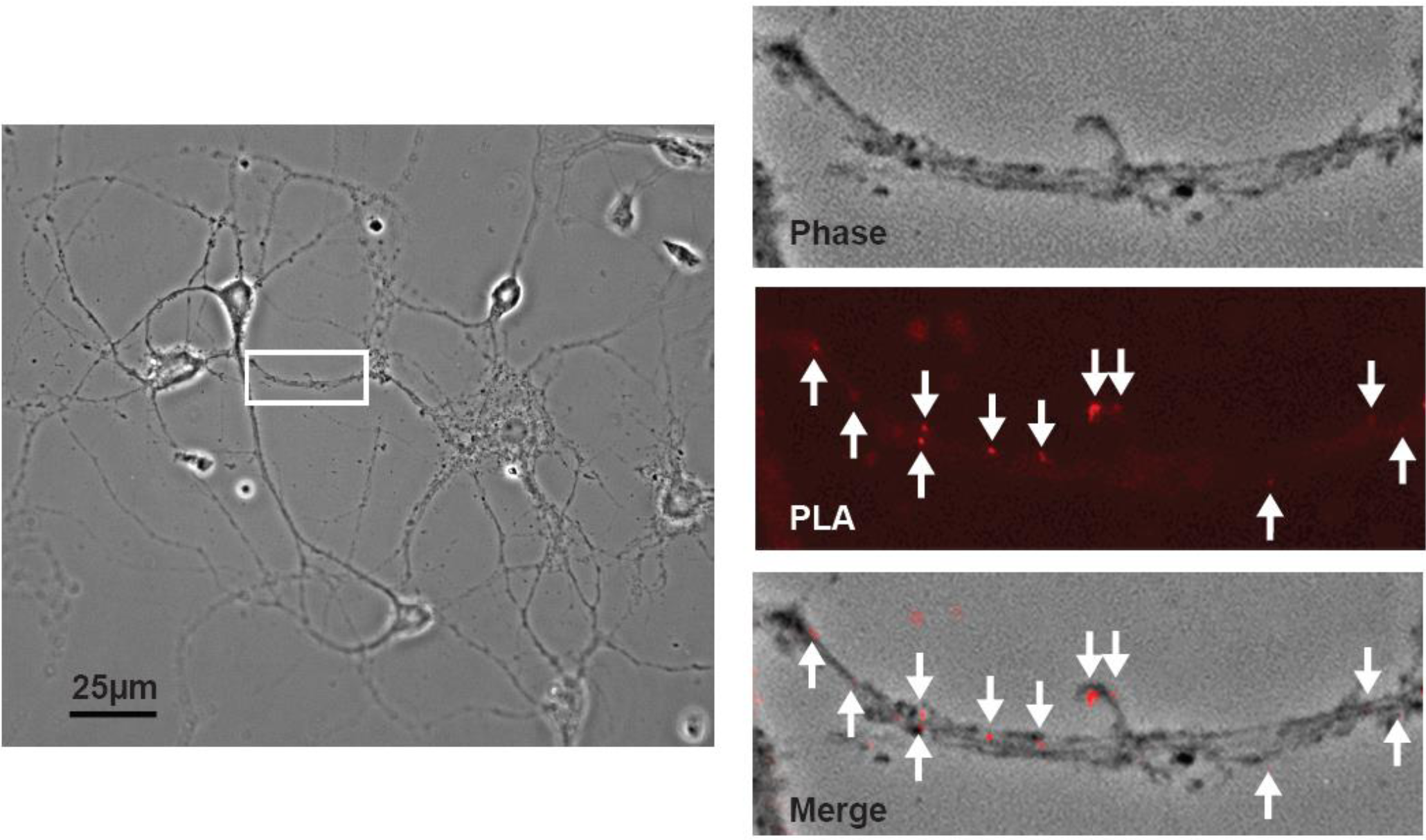
Demonstration of D1/NMDAR heteromers in cultured NAc neurons using a proximity ligation assay. A proximity ligation assay (PLA) was used to detect the dopamine D1 receptor and the obligatory NMDA receptor subunit GluN1 in sufficiently close proximity to represent heteromeric receptors. Left: Phase contrast image of co-cultured NAc and PFC neurons, with white rectangle identifying a NAc MSN process that is depicted at higher magnification in other panels. Right: PLA signals (red dots) indicate the presence of D1/NMDAR heteromers.

**Figure 7.**
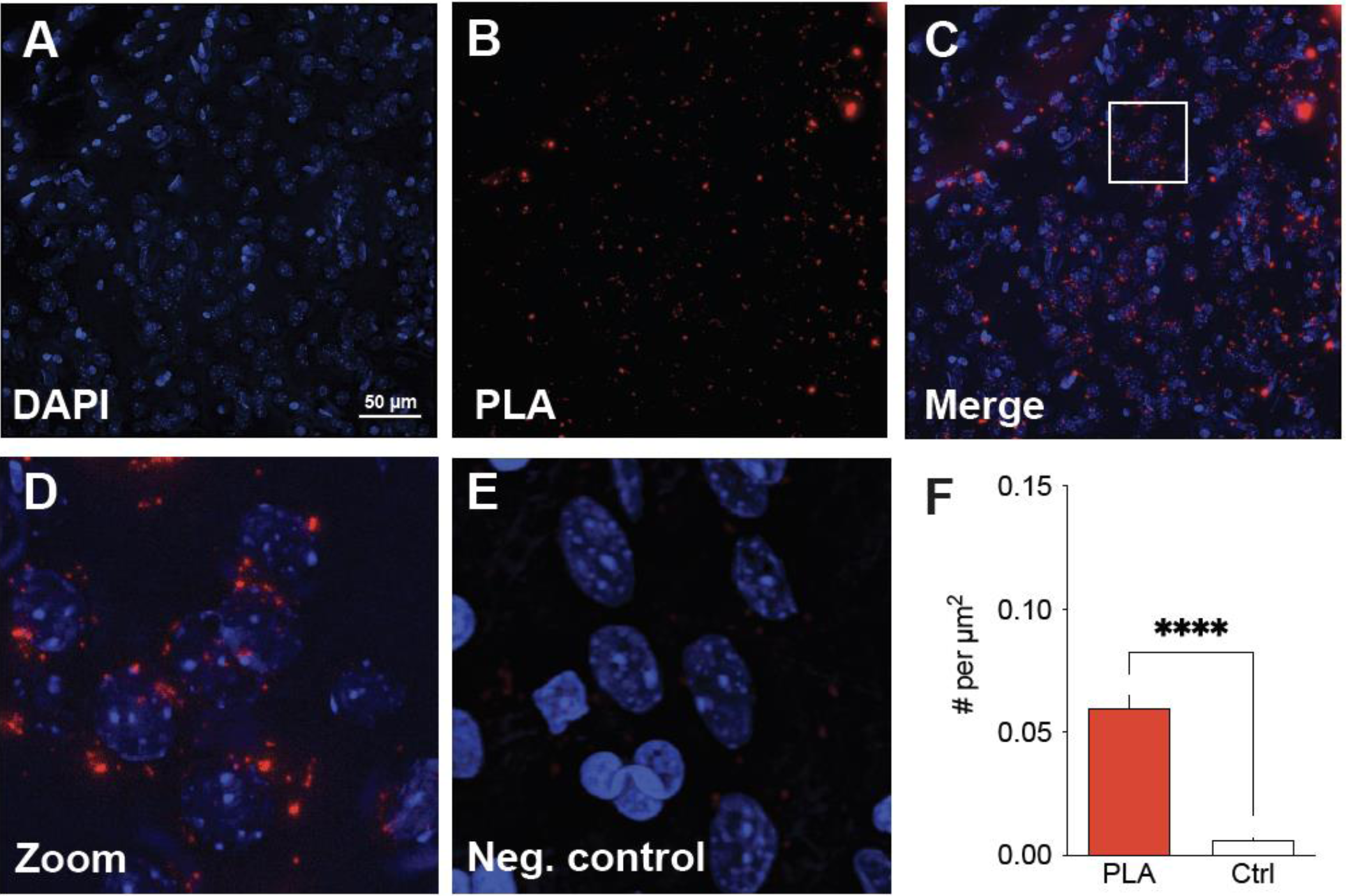
Evidence for the existence of D1/NMDAR heteromers in the adult rat NAc. A proximity ligation assay (PLA) was used in 30 µm coronal slices of the rat brain to detect the dopamine D1 receptor and the obligatory NMDAR subunit GluN1 in sufficiently close proximity to represent heteromeric receptors. (A-C): Representative images from the NAc show neuronal nuclei (stained by DAPI; A), PLA signals (red dots; B), and merged DAPI and PLA images demonstrating D1/NMDA heteromers in proximity to NAc cell bodies (C). (D) Higher magnification image of white box in panel C showing PLA labeling. E. High magnification negative control image showing absence of PLA labeling if the D1 receptor primary antibody is omitted. F. Significantly more puncta are detected in PLA experiments using D1 and GluN1 antibodies (e.g., panel D) compared to experiments in which the D1 antibody was omitted (e.g., panel E). Data in panel F are based on 12 images, obtained from a total of 6 slices from 3 male rats. Unpaired t-test, t(10) = 10.98, p < 0.0001.

To address the functional significance of D1R/NMDAR heteromers, additional co-cultures were co-incubated with AHA and bicuculline, plus either APV (50 µM), SCH23390 (10 µM), or both APV and SCH23390 (Fig. 8). APV alone eliminated the bicuculline-stimulated translation, confirming previous results [35]. SCH23390 alone also abolished bicuculline-stimulated translation, confirming results of our prior experiment (Fig. 2). Combining APV+SCH23390, however, did not result in any further reduction in the AHA signal, suggesting that antagonist occupation of D1Rs within the heteromers is sufficient to prevent their activation by glutamate. The alternative interpretation, namely that each receptor is independently capable of maximally reducing translation, is not consistent with the absence of any source of DA tone in the co-culture system.

**Figure 8.**
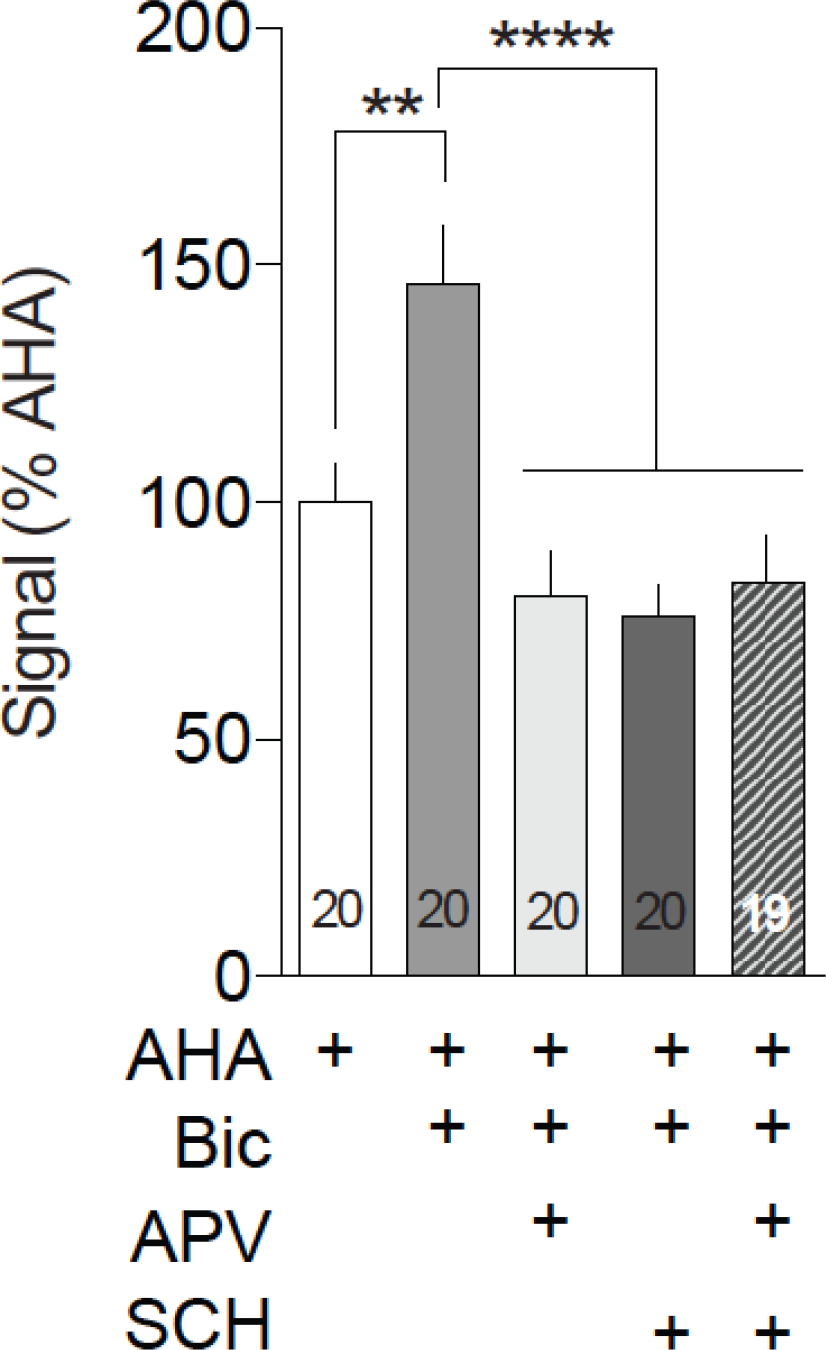
The effects of NMDA receptor and D1-class receptor blockade on bicuculline-stimulated NAc protein translation are not additive. Co-cultured NAc and PFC neurons were incubated with 1 mM AHA plus or minus the GABA_A_ receptor antagonist bicuculline (Bic; 20 µM) and/or the NMDA receptor antagonist APV (50 µm) or the dopamine D1/D5 receptor antagonist SCH23390 (SCH; 1 µm) for 2 h and tagged with 20 nM DBCO-Cy5. Bicuculline increased translation in the processes of NAc MSNs [F(4,94)=2.053, p<0.0001; Tukey’s post hoc p=0.006]. This effect was blocked by co-incubation with APV (APV vs Bic; Tukey’s post hoc p<0.0001) or SCH (SCH vs Bic; Tukey’s post hoc p<0.0001) and by a combination of APV and SCH (APV + SCH vs Bic; Tukey’s post hoc p<0.0001), but the effect of APV was not further impacted by co-incubation of APV with SCH (APV + SCH vs APV; Tukey’s post hoc p=0.9996; APV + SCH vs SCH; Tukey’s post hoc p=0.9839). **p<0.005, ****p<0.0001

## DISCUSSION

The goal of this study was to determine whether DA receptors modulate protein translation in NAc MSNs. We used a co-culture system in which PFC neurons are plated with NAc neurons in order to restore excitatory synapses onto the NAc MSNs. Newly synthesized proteins were identified using the FUNCAT technique, in which the non-canonical amino acid AHA incorporates into elongating peptides and can subsequently be labeled with Cy5 using click chemistry [37]. Levels of Cy5 fluorescence, our measure of protein translation, were then quantified using immunocytochemistry in processes of NAc MSNs. We examined basal translation and stimulated translation elicited by the GABA_A_ receptor antagonist bicuculline. We have shown previously that bicuculline-induced translation in MSNs in this co-culture system is glutamate-driven, since it can be eliminated by antagonists of NMDAR, AMPAR, or group I mGluRs [35], similar to previous findings on glutamate-driven protein translation in hippocampal cultures [57]. Our results show that neither D1-nor D2-class agonists affect basal translation, although a D2-class agonist reduced bicuculline-induced translation. Surprisingly, we also found that bicuculline-induced translation is blocked by either the NMDAR antagonist APV or the D1R antagonist SCH23390 (despite the fact that no DA-producing neurons exist in the co-cultures), with no additive effect observed when APV and SCH23390 are combined. We propose that our findings can be explained by D1R/NMDAR heteromers, which have been documented in other systems and which we now show are also present in NAc MSNs. Given the absence of DA in these cultures, our results suggest that glutamate activation of NMDARs within the D1R/NMDAR heteromers is sufficient to drive translation; however, antagonism of the D1R within these heteromers can prevent this activation.

### D1R/NMDAR heteromers

Multiple lines of evidence - including co-immunoprecipitation, GST-pull down, BRET, mass spectrometry, single-molecule imaging, and PLA assays - support the existence of D1R/NMDAR heteromers in hippocampal, prefrontal cortical and striatal neurons [58–66]. Interactions have been reported between the D1R C-terminal and the GluN1 C-terminal and between a distinct portion of the D1R C-terminal and the GluN2A C-terminal [59]. With the discovery of D1R/NMDAR heteromers, it became clear that some reported D1-NMDAR interactions reflect activation of signaling pathways downstream of D1Rs whereas others can be attributable to D1R/NMDAR heteromers (for review, see [67,68]). Probing the function of D1R/NMDAR heteromers has largely relied on peptides that selectively block either interaction. Disrupting the D1R-GluN2A association prevents D1R-mediated reduction of NMDAR function in HEK cells and hippocampal neurons [59]. On the other hand, disrupting the D1R-GluN1 association interferes with a broader array of events: neuroprotective effects of D1R stimulation [59]; D1R-mediated enhancement of excitatory synaptic transmission in hippocampal neurons, and a hippocampus-dependent spatial working memory task [69]; and, D1R-mediated enhancement of NMDAR currents and high frequency stimulation-induced LTP in D1R MSN, synergistic ERK activation in response to combined glutamate and D1-class agonist stimulation, and single injection-induced behavioral sensitization to cocaine [65]. It was recently shown that repeated cocaine exposure increased D1R/NMDAR heteromer formation and this was required for cocaine-induced potentiation of glutamate transmission in D1R MSN of NAc and behavioral sensitization [66]. Cocaine also influences D2R/NMDAR heteromerization as well as synaptic and behavioral responses to cocaine [66,70]. Formation vs dissociation of heteromers also influences the trafficking and subcellular localization of both D1Rs and NMDARs in striatal and hippocampal neurons [58,64,60,71]. Interestingly, some results suggest that D1R/NMDAR heteromers serve as a perisynaptic reserve pool that increases NMDAR synaptic content upon heteromer dissociation [64]. Overall, prior studies implicate D1R/NMDAR heteromers in a diverse array of processes. Our results, discussed below, suggest they also regulate protein translation.

### Regulation of protein translation by D1 and NMDARs in NAc MSNs

As noted above, PLA has previously been used to demonstrate D1R/NMDAR heteromers in MSN of dorsal striatum and NAc [65,66]. Our results provide the first evidence of D1R/NMDAR heteromers in NAc MSN of the rat. First, our immunocytochemical studies in cultured NAc MSNs revealed that a portion of D1R staining colocalizes with GluN1 staining. We then used PLA to demonstrate the existence of D1R/NMDAR heteromers both in cultured NAc MSNs and in NAc MSNs in brain slices from adult rats. We note that our PLA data do not reveal the identity of the NMDAR subunit with which D1Rs are interacting; the interaction could be with GluN1 itself, or with GluN2A present within a tetrameric GluN1-containing NMDAR. It would be interesting in the future to determine if NAc MSNs also express other types of oligomers containing both DA and glutamate receptors, including D2/GluN2B [70] and D1/Histamine H3/NMDAR complexes [72].

The original goal of the present study was to determine whether DA modulates protein translation in NAc MSNs. To examine potential modulation of basal protein translation, co-cultures were co-incubated for 2 h with AHA and either the D1-class agonist SKF81297 or the D2-class agonist quinpirole. Fig. 1A shows that neither DA agonist impacted translation relative to the AHA-only control group. This suggests that DA alone is not sufficient to drive protein translation in NAc MSNs in our co-culture system.

A disinhibitory effect of the GABA_A_ receptor antagonist bicuculline on excitatory neurotransmission in cultured cells is well established [73]. We have shown similar effects in our culture system [74,36]. More recently we demonstrated that bicuculline-induced disinhibition of excitatory transmission leads to increased protein translation [35], which we confirm here by demonstrating inhibition of bicuculline-stimulated translation with the NMDAR antagonist APV (Fig. 8). To assess the effect of DA transmission on bicuculline-stimulated protein translation, cells were co-incubated with bicuculline and either of the DA agonists (SKF81297 or quinpirole) during the AHA-labeling period. Bicuculline alone robustly increased translation, an effect that was reversed by quinpirole but not SKF81297 (Fig. 1B). This result suggests that D2-class receptors negatively regulate translation in NAc MSNs under conditions of high excitatory activity. This is the first demonstration that D2-class receptors modulate protein translation in MSNs. It might reflect inhibitory effects of D2-class receptors on excitatory synaptic transmission (e.g., [49]).

Interestingly bicuculline-stimulated translation was also prevented by co-incubation with the D1-class *antagonist* SCH23390 (Fig. 2A) but not the D2-class antagonist eticlopride (Fig. 2B), as well as being prevented by APV (Fig. 8). While this effect of SCH23390 is suggestive of endogenous DA tone at D1-class receptors on NAc MSNs, there are no DA neurons present in this co-culture system (Fig. 3). We propose an alternative explanation involving drug actions on D1R/NMDAR heteromers. Specifically, it is possible that bicuculline is activating translation by increasing glutamatergic tone at such heteromers, and that this response is abrogated through an allosteric interaction occurring when D1Rs in the heteromer are occupied by antagonist. We note that prior studies of D1R/NMDAR heteromer function have not examined an analogous situation, i.e., blocking the D1R in the absence of DA tone. This alternative explanation is consistent with our finding that APV and SCH23390 produce equivalent inhibition of biciculline-induced translation, and that no additive effect is observed when they are combined (Fig. 8). Furthermore, if NMDARs existing outside of D1R/NMDAR heteromers were sufficient to activate translation, there is no explanation for the effect of SCH23390. Our results provide the first evidence that functional consequences of D1R/NMDAR heteromer activation may include regulation of protein translation.

Whereas blocking the D1R in the absence of DA tone prevented bicuculline-stimulated translation (Fig. 2A and Fig. 8), combining a D1-class agonist with bicucuclline did not further enhance translation (Fig. 1B). This suggests that activation of NMDARs within D1R/NMDAR heteromers is sufficient to produce maximal activation of translation and that modulation of this function by the D1R will be important under conditions where a D1R antagonist has been administered. We have proposed above that the D1R antagonist prevents heteromer activation through an allosteric mechanism. It is also possible that D1R antagonists work by disrupting the D1R-NMDAR association. Arguing against this, a study using hippocampal tissue did not observe an effect of SCH 23390 on the ability of D1R antibody to immunoprecipitate GluN1 [60]. On the other hand, DA agonists can affect D1R-NMDAR physical interactions, although directionality is complex. For example, several lines of evidence suggest that D1R activation disrupts D1/NMDAR interactions in hippocampal neurons [64,59,69], but a D1-class agonist increased D1R/NMDAR heteromer formation, measured with PLA, in cultured striatal MSNs [65]. Future studies will use PLA to assess the effect of D1R stimulation and blockade on the number of D1R/NMDAR heteromers in MSNs in NAc slices and their effect on protein translation.

We did not distinguish between MSNs expressing D1Rs versus D2Rs because our culture system exhibits significantly less segregation of these receptors compared to MSNs in the adult rat (∼70% of MSNs in the co-cultures are D1R-positive and ∼80% are D2R-positive), most likely because our cultures are prepared from early postnatal brains [32].

### Relationship to prior findings in other experimental systems and brain regions

NMDAR modulation of protein translation depends on whether glutamate transmission is evoked or spontaneous, i.e., resulting from spontaneous fusion of a synaptic vesicle with the presynaptic membrane. In the hippocampus, NMDAR-mediated miniature synaptic events resulting from spontaneous transmission exert a suppressive effect on protein translation and thus NMDAR antagonists such as APV lead to increased translation [75–78]. We have similarly observed an APV-induced increase in protein translation in NAc tissue from adult drug-naïve rats [28] but not in NAc MSNs in co-culture [35], perhaps reflecting lower levels of spontaneous glutamate transmission in the co-cultures compared to intact NAc tissue, and in the NAc/PFC co-cultures versus hippocampal cultures (the latter are expected to contain a significantly higher portion of glutamate neurons). On the other hand, glutamate transmission evoked via bicuculline-induced disinhibition leads to an increase in protein translation in cultured NAc MSNs that is blocked by APV [35]. Likewise, glutamate transmission evoked with high potassium or glutamate agonists leads to an APV-sensitive increase in translation in processes of cultured hippocampal neurons [57]. Our results suggest a role for D1R/NMDAR heteromers in modulation of translation by evoked glutamate transmission.

Likewise, the effect of DA receptor stimulation on translation depends on experimental conditions, although here we are referring to distinctions between neurons studied under baseline versus stimulated conditions, and this is not equivalent to distinguishing spontaneous versus evoked glutamate transmission, as above. Our results show that DA agonists do not affect translation in cultured NAc MSNs studied under baseline conditions (Fig. 1A), whereas quinpirole but not SKF 81297 reversed bicuculline-stimulated protein translation (Fig. 1B). This contrasts with the hippocampus, where a D1-class agonist (SKF38393 or SKF81297; 25-100 µM) increased protein translation in the absence of other stimulation [15,16]. We used a lower concentration of D1-class agonist (SKF81297; 1 µM) but this is a saturating concentration (e.g., [60]) and we previously showed that this concentration was sufficient to modulate AMPAR trafficking in MSNs in NAc/PFC co-cultures [32] as well as NAc neurons cultured alone [33,34] and in other types of cultured neurons [49,79]. The timing of D1-class agonist treatment is probably not the explanation. We used a 2 h incubation time because this produces optimal AHA labeling in our culture system [35] and because we were concerned that restricting D1-class agonist exposure to only a portion of this incubation period might lead to masking of D1R-mediated effects. In hippocampal neurons, incubation with D1-class agonists for periods ranging from 15 min to 2.5 h increased protein translation [15,16]. Furthermore, it is clear that DA can modulate protein synthesis-dependent processes over longer time frames resembling our 2 h incubation. Thus, application of DA or a D1-class agonist to hippocampal slices produced a slowly developing potentiation of synaptic transmission that peaked 3-4 h after drug application and depended on protein translation [10,12,80]. Related to this is the well-established contribution of D1R-PKA signaling to the late protein synthesis-dependent stage of LTP [10,81,12–14].

### Conclusions

Our results show that DA does not affect protein translation under basal conditions in NAc MSNs in co-culture with PFC neurons, whereas D2-class receptors negatively regulate the enhanced translation driven by disinhibition of excitatory transmission with bicuculline. Most surprisingly, our results also point to a critical role for D1R/NMDAR heteromers in bicuculline-induced protein translation in NAc MSNs, with antagonist occupation of the D1R in the heteromer preventing NMDAR enhancement of translation. More work is required to further characterize this function of D1R/NMDAR heteromers and then determine why they might have privileged access, compared to either receptor alone, to regulation of protein translation. Finally, given the important role for protein translation in cocaine-induced plasticity [21–31], it will be interesting to determine if regulation of translation by D2 receptors and D1/NMDAR heteromers contributes to synaptic and behavioral effects of cocaine.

## Acknowledgements

This work is supported by grants DA015835 (MEW) and DA040414 (MTS) from the National Institute on Drug Abuse (United States). We thank Dr. Susan George for her generous gift of the D1 primary antibody. We thank Dr. Pablo Castillo and Hannah Monday for providing their protocol for FUNCAT in cultured neurons. We thank Dr. Amanda Wunsch for assistance with preparation of figures.

## Ethical approval

All applicable international, national, and/or institutional guidelines for the care and use of animals were followed. Specifically, all procedures performed in studies involving animals were in accordance with the ethical standards of the Institutional Animal Care and Use Committee of Rosalind Franklin University of Medicine and Science and North Central College.

## References

1. Sesack SR, Grace AA (2010) Cortico-Basal Ganglia reward network: microcircuitry. Neuropsychopharmacology 35 (1):27–47. doi:10.1038/npp.2009.93

2. Sheynikhovich D, Otani S, Bai J, Arleo A (2022) Long-term memory, synaptic plasticity and dopamine in rodent medial prefrontal cortex: Role in executive functions. Front Behav Neurosci 16:1068271. doi:10.3389/fnbeh.2022.1068271

3. Sippy T, Tritsch NX (2023) Unraveling the dynamics of dopamine release and its actions on target cells. Trends Neurosci 46 (3):228–239. doi:10.1016/j.tins.2022.12.005

4. Tritsch NX, Sabatini BL (2012) Dopaminergic modulation of synaptic transmission in cortex and striatum. Neuron 76 (1):33–50. doi:10.1016/j.neuron.2012.09.023

5. Tsetsenis T, Broussard JI, Dani JA (2022) Dopaminergic regulation of hippocampal plasticity, learning, and memory. Front Behav Neurosci 16:1092420. doi:10.3389/fnbeh.2022.1092420

6. Abraham WC, Williams JM (2008) LTP maintenance and its protein synthesis-dependence. Neurobiology of learning and memory 89 (3):260–268. doi:10.1016/j.nlm.2007.10.001

7. Zukin RS, Richter JD, Bagni C (2009) Signals, synapses, and synthesis: how new proteins control plasticity. Frontiers in neural circuits 3:14. doi:10.3389/neuro.04.014.2009

8. Buffington SA, Huang W, Costa-Mattioli M (2014) Translational control in synaptic plasticity and cognitive dysfunction. Annual review of neuroscience 37:17–38. doi:10.1146/annurev-neuro-071013-014100

9. Sutton MA, Schuman EM (2006) Dendritic protein synthesis, synaptic plasticity, and memory. Cell 127 (1):49–58. doi:10.1016/j.cell.2006.09.014

10. Frey U, Huang YY, Kandel ER (1993) Effects of cAMP simulate a late stage of LTP in hippocampal CA1 neurons. Science 260 (5114):1661–1664

11. Frey S, Schwiegert C, Krug M, Lossner B (1991) Long-term potentiation induced changes in protein synthesis of hippocampal subfields of freely moving rats: time-course. Biomed Biochim Acta 50 (12):1231–1240

12. Huang YY, Kandel ER (1995) D1/D5 receptor agonists induce a protein synthesis-dependent late potentiation in the CA1 region of the hippocampus. Proceedings of the National Academy of Sciences of the United States of America 92 (7):2446–2450

13. Matthies H, Becker A, Schroeder H, Kraus J, Hollt V, Krug M (1997) Dopamine D1-deficient mutant mice do not express the late phase of hippocampal long-term potentiation. Neuroreport 8 (16):3533–3535

14. Swanson-Park JL, Coussens CM, Mason-Parker SE, Raymond CR, Hargreaves EL, Dragunow M, Cohen AS, Abraham WC (1999) A double dissociation within the hippocampus of dopamine D1/D5 receptor and beta-adrenergic receptor contributions to the persistence of long-term potentiation. Neuroscience 92 (2):485–497

15. Smith WB, Starck SR, Roberts RW, Schuman EM (2005) Dopaminergic stimulation of local protein synthesis enhances surface expression of GluR1 and synaptic transmission in hippocampal neurons. Neuron 45 (5):765–779. doi:10.1016/j.neuron.2005.01.015

16. Hodas JJ, Nehring A, Hoche N, Sweredoski MJ, Pielot R, Hess S, Tirrell DA, Dieterich DC, Schuman EM (2012) Dopaminergic modulation of the hippocampal neuropil proteome identified by bioorthogonal noncanonical amino acid tagging (BONCAT). Proteomics 12 (15-16):2464–2476. doi:10.1002/pmic.201200112

17. Koob GF, Volkow ND (2010) Neurocircuitry of addiction. Neuropsychopharmacology : official publication of the American College of Neuropsychopharmacology 35 (1):217–238. doi:10.1038/npp.2009.110

18. Scofield MD, Heinsbroek JA, Gipson CD, Kupchik YM, Spencer S, Smith AC, Roberts-Wolfe D, Kalivas PW (2016) The Nucleus Accumbens: Mechanisms of Addiction across Drug Classes Reflect the Importance of Glutamate Homeostasis. Pharmacol Rev 68 (3):816–871. doi:10.1124/pr.116.012484

19. Luscher C (2016) The Emergence of a Circuit Model for Addiction. Annual review of neuroscience 39:257–276. doi:10.1146/annurev-neuro-070815-013920

20. Wolf ME (2016) Synaptic mechanisms underlying persistent cocaine craving. Nat Rev Neurosci 17 (6):351–365. doi:10.1038/nrn.2016.39

21. Huang W, Placzek AN, Viana Di Prisco G, Khatiwada S, Sidrauski C, Krnjevic K, Walter P, Dani JA, Costa-Mattioli M (2016) Translational control by eIF2alpha phosphorylation regulates vulnerability to the synaptic and behavioral effects of cocaine. Elife 5. doi:10.7554/eLife.12052

22. Neasta J, Barak S, Hamida SB, Ron D (2014) mTOR complex 1: a key player in neuroadaptations induced by drugs of abuse. Journal of neurochemistry 130 (2):172–184. doi:10.1111/jnc.12725

23. Scheyer AF, Wolf ME, Tseng KY (2014) A protein synthesis-dependent mechanism sustains calcium-permeable AMPA receptor transmission in nucleus accumbens synapses during withdrawal from cocaine self-administration. J Neurosci 34 (8):3095–3100. doi:10.1523/JNEUROSCI.4940-13.2014

24. Dayas CV, Smith DW, Dunkley PR (2012) An emerging role for the Mammalian target of rapamycin in “pathological” protein translation: relevance to cocaine addiction. Front Pharmacol 3:13. doi:10.3389/fphar.2012.00013

25. Placzek AN, Molfese DL, Khatiwada S, Viana Di Prisco G, Huang W, Sidrauski C, Krnjevic K, Amos CL, Ray R, Dani JA, Walter P, Salas R, Costa-Mattioli M (2016) Translational control of nicotine-evoked synaptic potentiation in mice and neuronal responses in human smokers by eIF2alpha. Elife 5. doi:10.7554/eLife.12056

26. Placzek AN, Prisco GV, Khatiwada S, Sgritta M, Huang W, Krnjevic K, Kaufman RJ, Dani JA, Walter P, Costa-Mattioli M (2016) eIF2alpha-mediated translational control regulates the persistence of cocaine-induced LTP in midbrain dopamine neurons. Elife 5. doi:10.7554/eLife.17517

27. Beckley JT, Laguesse S, Phamluong K, Morisot N, Wegner SA, Ron D (2016) The First Alcohol Drink Triggers mTORC1-Dependent Synaptic Plasticity in Nucleus Accumbens Dopamine D1 Receptor Neurons. The Journal of neuroscience : the official journal of the Society for Neuroscience 36 (3):701–713. doi:10.1523/JNEUROSCI.2254-15.2016

28. Stefanik MT, Milovanovic M, Werner CT, Spainhour JCG, Wolf ME (2018) Withdrawal From Cocaine Self-administration Alters the Regulation of Protein Translation in the Nucleus Accumbens. Biol Psychiatry 84 (3):223–232. doi:10.1016/j.biopsych.2018.02.012

29. Werner CT, Stefanik MT, Milovanovic M, Caccamise A, Wolf ME (2018) Protein Translation in the Nucleus Accumbens Is Dysregulated during Cocaine Withdrawal and Required for Expression of Incubation of Cocaine Craving. J Neurosci 38 (11):2683–2697. doi:10.1523/JNEUROSCI.2412-17.2018

30. Laguesse S, Ron D (2019) Protein Translation and Psychiatric Disorders. Neuroscientist:1073858419853236. doi:10.1177/1073858419853236

31. Kawa AB, Hwang EK, Funke JR, Zhou H, Costa-Mattioli M, Wolf ME (2022) Positive Allosteric Modulation of mGlu(1) Reverses Cocaine-Induced Behavioral and Synaptic Plasticity Through the Integrated Stress Response and Oligophrenin-1. Biol Psychiatry 92 (11):871–879. doi:10.1016/j.biopsych.2022.05.008

32. Sun X, Milovanovic M, Zhao Y, Wolf ME (2008) Acute and chronic dopamine receptor stimulation modulates AMPA receptor trafficking in nucleus accumbens neurons cocultured with prefrontal cortex neurons. The Journal of neuroscience : the official journal of the Society for Neuroscience 28 (16):4216–4230. doi:10.1523/JNEUROSCI.0258-08.2008

33. Chao SZ, Ariano MA, Peterson DA, Wolf ME (2002) D1 dopamine receptor stimulation increases GluR1 surface expression in nucleus accumbens neurons. Journal of neurochemistry 83 (3):704–712

34. Mangiavacchi S, Wolf ME (2004) D1 dopamine receptor stimulation increases the rate of AMPA receptor insertion onto the surface of cultured nucleus accumbens neurons through a pathway dependent on protein kinase A. Journal of neurochemistry 88 (5):1261–1271

35. Stefanik MT, Sakas C, Lee D, Wolf ME (2018) Ionotropic and metabotropic glutamate receptors regulate protein translation in co-cultured nucleus accumbens and prefrontal cortex neurons. Neuropharmacology 140:62–75. doi:10.1016/j.neuropharm.2018.05.032

36. Sun X, Wolf ME (2009) Nucleus accumbens neurons exhibit synaptic scaling that is occluded by repeated dopamine pre-exposure. Eur J Neurosci 30 (4):539–550. doi:10.1111/j.1460-9568.2009.06852.x

37. Dieterich DC, Hodas JJ, Gouzer G, Shadrin IY, Ngo JT, Triller A, Tirrell DA, Schuman EM (2010) In situ visualization and dynamics of newly synthesized proteins in rat hippocampal neurons. Nature neuroscience 13 (7):897–905. doi:10.1038/nn.2580

38. Hinz FI, Dieterich DC, Tirrell DA, Schuman EM (2012) Non-canonical amino acid labeling in vivo to visualize and affinity purify newly synthesized proteins in larval zebrafish. ACS Chem Neurosci 3 (1):40–49. doi:10.1021/cn2000876

39. Tom Dieck S, Muller A, Nehring A, Hinz FI, Bartnik I, Schuman EM, Dieterich DC (2012) Metabolic labeling with noncanonical amino acids and visualization by chemoselective fluorescent tagging. Curr Protoc Cell Biol Chapter 7:Unit7 11. doi:10.1002/0471143030.cb0711s56

40. Gomes I, Sierra S, Devi LA (2016) Detection of Receptor Heteromerization Using In Situ Proximity Ligation Assay. Curr Protoc Pharmacol 75:2 16 11-12 16 31. doi:10.1002/cpph.15

41. Soderberg O, Gullberg M, Jarvius M, Ridderstrale K, Leuchowius KJ, Jarvius J, Wester K, Hydbring P, Bahram F, Larsson LG, Landegren U (2006) Direct observation of individual endogenous protein complexes in situ by proximity ligation. Nat Methods 3 (12):995–1000. doi:10.1038/nmeth947

42. Soderberg O, Leuchowius KJ, Gullberg M, Jarvius M, Weibrecht I, Larsson LG, Landegren U (2008) Characterizing proteins and their interactions in cells and tissues using the in situ proximity ligation assay. Methods 45 (3):227–232. doi:10.1016/j.ymeth.2008.06.014

43. Tom Dieck S, Kochen L, Hanus C, Heumuller M, Bartnik I, Nassim-Assir B, Merk K, Mosler T, Garg S, Bunse S, Tirrell DA, Schuman EM (2015) Direct visualization of newly synthesized target proteins in situ. Nature methods 12 (5):411–414. doi:10.1038/nmeth.3319

44. Hinz FI, Dieterich DC, Schuman EM (2013) Teaching old NCATs new tricks: using non-canonical amino acid tagging to study neuronal plasticity. Curr Opin Chem Biol 17 (5):738–746. doi:10.1016/j.cbpa.2013.07.021

45. Bagert JD, Xie YJ, Sweredoski MJ, Qi Y, Hess S, Schuman EM, Tirrell DA (2014) Quantitative, time-resolved proteomic analysis by combining bioorthogonal noncanonical amino acid tagging and pulsed stable isotope labeling by amino acids in cell culture. Mol Cell Proteomics 13 (5):1352–1358. doi:10.1074/mcp.M113.031914

46. Sletten EM, Bertozzi CR (2009) Bioorthogonal chemistry: fishing for selectivity in a sea of functionality. Angew Chem Int Ed Engl 48 (38):6974–6998. doi:10.1002/anie.200900942

47. Biever A, Valjent E, Puighermanal E (2015) Ribosomal Protein S6 Phosphorylation in the Nervous System: From Regulation to Function. Frontiers in molecular neuroscience 8:75. doi:10.3389/fnmol.2015.00075

48. Graber TE, Hebert-Seropian S, Khoutorsky A, David A, Yewdell JW, Lacaille JC, Sossin WS (2013) Reactivation of stalled polyribosomes in synaptic plasticity. Proceedings of the National Academy of Sciences of the United States of America 110 (40):16205–16210. doi:10.1073/pnas.1307747110

49. Sun X, Zhao Y, Wolf ME (2005) Dopamine receptor stimulation modulates AMPA receptor synaptic insertion in prefrontal cortex neurons. The Journal of neuroscience : the official journal of the Society for Neuroscience 25 (32):7342–7351. doi:10.1523/JNEUROSCI.4603-04.2005

50. Lee SP, So CH, Rashid AJ, Varghese G, Cheng R, Lanca AJ, O’Dowd BF, George SR (2004) Dopamine D1 and D2 receptor Co-activation generates a novel phospholipase C-mediated calcium signal. J Biol Chem 279 (34):35671–35678. doi:10.1074/jbc.M401923200

51. Hasbi A, Perreault ML, Shen MYF, Fan T, Nguyen T, Alijaniaram M, Banasikowski TJ, Grace AA, O’Dowd BF, Fletcher PJ, George SR (2017) Activation of Dopamine D1-D2 Receptor Complex Attenuates Cocaine Reward and Reinstatement of Cocaine-Seeking through Inhibition of DARPP-32, ERK, and DeltaFosB. Front Pharmacol 8:924. doi:10.3389/fphar.2017.00924

52. Trifilieff P, Rives ML, Urizar E, Piskorowski RA, Vishwasrao HD, Castrillon J, Schmauss C, Slattman M, Gullberg M, Javitch JA (2011) Detection of antigen interactions ex vivo by proximity ligation assay: endogenous dopamine D2-adenosine A2A receptor complexes in the striatum. Biotechniques 51 (2):111–118. doi:10.2144/000113719

53. Lu W, Shi Y, Jackson AC, Bjorgan K, During MJ, Sprengel R, Seeburg PH, Nicoll RA (2009) Subunit composition of synaptic AMPA receptors revealed by a single-cell genetic approach. Neuron 62 (2):254–268. doi:10.1016/j.neuron.2009.02.027

54. Reimers JM, Milovanovic M, Wolf ME (2011) Quantitative analysis of AMPA receptor subunit composition in addiction-related brain regions. Brain research 1367:223–233. doi:10.1016/j.brainres.2010.10.016

55. Wenthold RJ, Petralia RS, Blahos J, II, Niedzielski AS (1996) Evidence for multiple AMPA receptor complexes in hippocampal CA1/CA2 neurons. The Journal of neuroscience : the official journal of the Society for Neuroscience 16 (6):1982–1989

56. Conrad KL, Tseng KY, Uejima JL, Reimers JM, Heng LJ, Shaham Y, Marinelli M, Wolf ME (2008) Formation of accumbens GluR2-lacking AMPA receptors mediates incubation of cocaine craving. Nature 454 (7200):118–121. doi:10.1038/nature06995

57. Gong R, Park CS, Abbassi NR, Tang SJ (2006) Roles of glutamate receptors and the mammalian target of rapamycin (mTOR) signaling pathway in activity-dependent dendritic protein synthesis in hippocampal neurons. J Biol Chem 281 (27):18802–18815. doi:10.1074/jbc.M512524200

58. Fiorentini C, Gardoni F, Spano P, Di Luca M, Missale C (2003) Regulation of dopamine D1 receptor trafficking and desensitization by oligomerization with glutamate N-methyl-D-aspartate receptors. J Biol Chem 278 (22):20196–20202. doi:10.1074/jbc.M213140200

59. Lee FJ, Xue S, Pei L, Vukusic B, Chery N, Wang Y, Wang YT, Niznik HB, Yu XM, Liu F (2002) Dual regulation of NMDA receptor functions by direct protein-protein interactions with the dopamine D1 receptor. Cell 111 (2):219–230

60. Pei L, Lee FJ, Moszczynska A, Vukusic B, Liu F (2004) Regulation of dopamine D1 receptor function by physical interaction with the NMDA receptors. The Journal of neuroscience : the official journal of the Society for Neuroscience 24 (5):1149–1158. doi:10.1523/JNEUROSCI.3922-03.2004

61. Woods AS, Ciruela F, Fuxe K, Agnati LF, Lluis C, Franco R, Ferre S (2005) Role of electrostatic interaction in receptor-receptor heteromerization. J Mol Neurosci 26 (2-3):125–132. doi:10.1385/JMN:26:2-3:125

62. Woods AS, Ferre S (2005) Amazing stability of the arginine-phosphate electrostatic interaction. J Proteome Res 4 (4):1397–1402. doi:10.1021/pr050077s

63. Kruse MS, Premont J, Krebs MO, Jay TM (2009) Interaction of dopamine D1 with NMDA NR1 receptors in rat prefrontal cortex. Eur Neuropsychopharmacol 19 (4):296–304. doi:10.1016/j.euroneuro.2008.12.006

64. Ladepeche L, Dupuis JP, Bouchet D, Doudnikoff E, Yang L, Campagne Y, Bezard E, Hosy E, Groc L (2013) Single-molecule imaging of the functional crosstalk between surface NMDA and dopamine D1 receptors. Proceedings of the National Academy of Sciences of the United States of America 110 (44):18005–18010. doi:10.1073/pnas.1310145110

65. Cahill E, Pascoli V, Trifilieff P, Savoldi D, Kappes V, Luscher C, Caboche J, Vanhoutte P (2014) D1R/GluN1 complexes in the striatum integrate dopamine and glutamate signalling to control synaptic plasticity and cocaine-induced responses. Mol Psychiatry 19 (12):1295–1304. doi:10.1038/mp.2014.73

66. Andrianarivelo A, Saint-Jour E, Pousinha P, Fernandez SP, Petitbon A, De Smedt-Peyrusse V, Heck N, Ortiz V, Allichon MC, Kappes V, Betuing S, Walle R, Zhu Y, Josephine C, Bemelmans AP, Turecki G, Mechawar N, Javitch JA, Caboche J, Trifilieff P, Barik J, Vanhoutte P (2021) Disrupting D1-NMDA or D2-NMDA receptor heteromerization prevents cocaine’s rewarding effects but preserves natural reward processing. Sci Adv 7 (43):eabg5970. doi:10.1126/sciadv.abg5970

67. Cepeda C, Levine MS (2006) Where do you think you are going? The NMDA-D1 receptor trap. Sci STKE 2006 (333):pe20. doi:10.1126/stke.3332006pe20

68. Wang M, Wong AH, Liu F (2012) Interactions between NMDA and dopamine receptors: a potential therapeutic target. Brain research 1476:154–163. doi:10.1016/j.brainres.2012.03.029

69. Nai Q, Li S, Wang SH, Liu J, Lee FJ, Frankland PW, Liu F (2010) Uncoupling the D1-N-methyl-D-aspartate (NMDA) receptor complex promotes NMDA-dependent long-term potentiation and working memory. Biol Psychiatry 67 (3):246–254. doi:10.1016/j.biopsych.2009.08.011

70. Liu XY, Chu XP, Mao LM, Wang M, Lan HX, Li MH, Zhang GC, Parelkar NK, Fibuch EE, Haines M, Neve KA, Liu F, Xiong ZG, Wang JQ (2006) Modulation of D2R-NR2B interactions in response to cocaine. Neuron 52 (5):897–909. doi:10.1016/j.neuron.2006.10.011

71. Scott L, Zelenin S, Malmersjo S, Kowalewski JM, Markus EZ, Nairn AC, Greengard P, Brismar H, Aperia A (2006) Allosteric changes of the NMDA receptor trap diffusible dopamine 1 receptors in spines. Proceedings of the National Academy of Sciences of the United States of America 103 (3):762–767. doi:10.1073/pnas.0505557103

72. Rodriguez-Ruiz M, Moreno E, Moreno-Delgado D, Navarro G, Mallol J, Cortes A, Lluis C, Canela EI, Casado V, McCormick PJ, Franco R (2017) Heteroreceptor Complexes Formed by Dopamine D1, Histamine H3, and N-Methyl-D-Aspartate Glutamate Receptors as Targets to Prevent Neuronal Death in Alzheimer’s Disease. Molecular neurobiology 54 (6):4537-4550. doi:10.1007/s12035-016-9995-y

73. Turrigiano GG, Nelson SB (2004) Homeostatic plasticity in the developing nervous system. Nature reviews Neuroscience 5 (2):97–107. doi:10.1038/nrn1327

74. Reimers JM, Loweth JA, Wolf ME (2014) BDNF contributes to both rapid and homeostatic alterations in AMPA receptor surface expression in nucleus accumbens medium spiny neurons. The European journal of neuroscience 39 (7):1159–1169. doi:10.1111/ejn.12422

75. Kavalali ET, Monteggia LM (2012) Synaptic mechanisms underlying rapid antidepressant action of ketamine. Am J Psychiatry 169 (11):1150–1156. doi:10.1176/appi.ajp.2012.12040531

76. Sutton MA, Ito HT, Cressy P, Kempf C, Woo JC, Schuman EM (2006) Miniature neurotransmission stabilizes synaptic function via tonic suppression of local dendritic protein synthesis. Cell 125 (4):785–799. doi:10.1016/j.cell.2006.03.040

77. Sutton MA, Wall NR, Aakalu GN, Schuman EM (2004) Regulation of dendritic protein synthesis by miniature synaptic events. Science 304 (5679):1979–1983. doi:10.1126/science.1096202

78. Nosyreva E, Szabla K, Autry AE, Ryazanov AG, Monteggia LM, Kavalali ET (2013) Acute suppression of spontaneous neurotransmission drives synaptic potentiation. J Neurosci 33 (16):6990–7002. doi:10.1523/JNEUROSCI.4998-12.2013

79. Gao C, Sun X, Wolf ME (2006) Activation of D1 dopamine receptors increases surface expression of AMPA receptors and facilitates their synaptic incorporation in cultured hippocampal neurons. Journal of neurochemistry 98 (5):1664–1677. doi:10.1111/j.1471-4159.2006.03999.x

80. Gribkoff VK, Ashe JH (1984) Modulation by dopamine of population responses and cell membrane properties of hippocampal CA1 neurons in vitro. Brain research 292 (2):327–338

81. Frey U, Matthies H, Reymann KG, Matthies H (1991) The effect of dopaminergic D1 receptor blockade during tetanization on the expression of long-term potentiation in the rat CA1 region in vitro. Neuroscience letters 129 (1):111–114

